# Inhibition of histone deacetylase 1 or 2 reduces microglia activation through a gene expression independent mechanism

**DOI:** 10.1101/107649

**Authors:** Benjamin S. Durham, Ronald Grigg, Ian C. Wood

## Abstract

Histone deacetylase (HDAC) inhibitors prevent neural cell death in *in vivo* models of cerebral ischaemia, brain injury and neurodegenerative disease. One mechanism by which HDAC inhibitors may do this is by suppressing the excessive inflammatory response of chronically activated microglia. However, the molecular mechanisms underlying this anti-inflammatory effect and the specific HDAC responsible are not fully understood. Recent data from *in vivo* rodent studies has shown that inhibition of class I HDACs suppresses neuroinflammation and is neuroprotective. In our study we have identified that selective HDAC inhibition with inhibitors apicidin, MS-275 or MI-192, or specific knockdown of HDAC1 or 2 using siRNA, suppresses the expression of cytokines interleukin-6 (IL-6) and tumour necrosis factor-alpha (TNF-α) in BV2 murine microglia activated with lipopolysaccharide (LPS). Furthermore, we found that in the absence of HDAC1, HDAC2 is upregulated and these increased levels are compensatory, suggesting these two HDACs have redundancy in regulating the inflammatory response of microglia. Investigating the possible underlying anti-inflammatory mechanisms suggests an increase in protein expression is not important. Taken together, this study supports the idea that inhibitors selective towards HDAC1 or HDAC2, may be therapeutically useful for targeting neuroinflammation in brain injuries and neurodegenerative disease.

**Significance Statement:** The number of patients suffering a stroke or a neurodegenerative disease, such as Alzheimer’s is increasing These conditions are severely debilitating and are leading causes of mortality, with neural cell death and loss of brain tissue being a major feature. A number of mechanisms contribute to neuronal death, including inflammation in the brain, but we still lack clinical therapies to inhibit this. The work presented here provides further insight into potential molecular therapeutic targets called histone deacetylases (HDACs), which are thought to contribute to neural cell death by promoting inflammation. We show that down regulation of HDAC1 and 2 is sufficient to reduce this inflammatory response. Our findings have clinical relevance because they identify HDAC1 and 2 as promising targets for therapy.

## INTRODUCTION

Microglia are the innate immune cells of the brain that are responsible for the excessive and chronic neuroinflammatory response known to contribute to the pathogenesis of brain injury and disease (Block et al., 2007; Glass et al., 2010). Inhibitors of histone deacetylases (HDACs) 59reduce the inflammatory response of isolated microglia to stimulants such as lipopolysaccharide (LPS) (Suuronen et al., 2003; Peng et al., 2005; Chen et al., 2007; Faraco et al., 2009; Suh et al., 2010; Kannan et al., 2013). When delivered *in vivo*, HDAC inhibitors (HDACi) reduce neuroinflammation, promote neuroprotection and improve functional outcomes in models of cerebral ischaemia (Kim et al., 2007; Sinn et al., 2007; Xuan et al., 2012; Kim and Chuang, 2014), traumatic brain injury (Zhang et al., 2008; Shein and Shohami, 2011) and encephalomyelitis (Camelo et al., 2005; Zhang et al., 2010). Therapies for treating brain injury and disease are lacking, but these studies highlight a role for HDACs in neuroinflammation and suggest they are appropriate targets to inhibit.

The mechanism by which HDAC inhibition is anti-inflammatory is not understood, but we know HDACs remove acetyl groups from lysine residues on proteins including, histones (Strahl and Allis, 2000), enzymes and transcription factors (Glozak et al., 2005; Yao and Yang, 2011). As a consequence, deacetylation of histones promotes a compact chromatin structure and reduces gene expression, and deacetylation of specific lysine residues on transcription factors can modulate their activity (Gu and Roeder, 1997; Boyes et al., 1998). The identity of which acetylated proteins are responsible for the anti-inflammatory responses observed when using HDAC inhibitors is unclear.

There are 18 mammalian HDAC isoforms (class I HDACs (1, 2, 3 and 8), class II HDACs (4-7, 9 and 10), class III sirtuins (1-7) and the class IV HDAC11) and the majority of studies to date have focused on using non-selective HDAC inhibitors such as suberoylanilide hydroxamic acid (SAHA), trichostatin-A (TSA) and valproic acid (VPA) which inhibit the majority of these isoforms. As a result, we do not know which HDACs modulate the microglial inflammatory response and how inhibition of these leads to the acetylation of specific proteins to reduce neuroinflammation. Recent studies have begun to address these questions and have shown that selective inhibition of class I HDACs 1, 2 and 3 with the HDAC inhibitor MS-275 (Hu et al., 2003; Simonini et al., 2006; Beckers et al., 2007; Khan et al.,2008) can reduce neuroinflammation in a mouse model of Alzheimer’s disease (Zhang and Schluesener, 2013) and the inflammatory response of macrophages to the inflammatory stimulant LPS (Jeong et al., 2014).

In our study, we have used selective HDAC inhibitors and siRNA mediated knockdown to identify HDAC1 and HDAC2 as the key HDACs involved in the neuroinflammatory response of microglia. We show that selective class I HDAC inhibitors and siRNA to specifically knockdown HDAC1 and 2, both suppressed the expression of cytokines in BV2 murine microglia. Knockdown of HDAC1 alone resulted in a compensatory increase in the levels of HDAC2 and did not suppress cytokine expression, showing these two enzymes have redundancy in the neuroinflammatory response. We show that the HDACi are effective in the absence of new protein synthesis suggesting that the anti-inflammatory mechanism of HDACi does not involve increased protein expression. This identification suggests that HDAC 100 selective inhibitors may be therapeutically useful for targeting microglia and neuroinflammation, in brain injury and disease by modulating the acetylation levels and function of non-histone protein(s).

## MATERIALS & METHODS

### CELL CULTURE

BV2 murine microglia were cultured in Dulbecco’s Modified Eagle’s Medium (DMEM) high glucose AQmedia™ (Sigma-Aldrich) supplemented with 10% v/v foetal bovine serum (FBS, PAA Cell Culture Company) and 100U penicillin/100 μg streptomycin (Sigma-Aldrich). Cells were seeded into 6-well plates at either 350,000 or 500,000 cells/well and 24-well plates at 175,000 cells/well. The cells were cultured for 24 hours before treatment. BV2 cells were treated with vehicle control, or HDAC inhibitor with or without 500 ng/mL lipopolysaccharide (LPS, Sigma) and cells harvested after 6 or 24 hours. Inhibitors used were; apicidin (Sigma-Aldrich), MI192 (School of Chemistry, University of Leeds), MS-275 (Cayman Chemicals), SAHA (Cayman Chemicals) all dissolved in DMSO and valproic acid (VPA, Sigma-Aldrich) dissolved in phosphate buffered saline (PBS, Oxoid). For pre-treatment experiments, BV2 cells were treated with vehicle control or appropriate drugs for 24 hours before the addition of 500 ng/mL LPS for a further 6 hours.

### CELL TRANSFECTION

BV2 microglia seeded into 6-well plates were washed with 1 mL PBS and maintained in 1 mL Opti-MEM® (Gibco) throughout the transfection procedure. Cells were transfected with 50 pmoles of Silencer® Select Negative Control siRNA (Ambion) or Silencer® Select Pre-designed siRNA targeted against HDAC1 (id: s119557, Ambion) or HDAC2 (id: s67417, Ambion) as follows. For each well, 3 μL of Lipofectamine™ 2000 (Invitrogen) was dissolved in 100 μL of Opti-MEM® and 1 μL of 50 μM siRNA was dissolved in 100 μL of Opti-Mem®, these were incubated for 5 minutes before combining and incubating for a further 20 minutes at room temperature. Afterwards, 200 μL of this mix was added to the well, followed by incubation at 37°C in a humid atmosphere with 5% CO_2_ for 4 hours. This was then removed and the cells were cultured in 3 mL DMEM high glucose AQmedia™ supplemented with 1%v/v FBS and 100U penicillin/100 μg streptomycin for 24 hours. Medium was then changed and the cells were cultured for a further 24 hours before treating with 500 ng/mL LPS for 6 hours.

### WHOLE CELL PROTEIN EXTRACTION

BV2 cells seeded into 6-well plates were washed with 1 mL PBS and scraped into 250 μL ice-cold RIPA Buffer [10 mM Tris-HCL pH 8.0, 1 mM EDTA, 1% Triton X-100, 0.1% sodium deoxycholate, 0.1% sodium dodecyl sulfate (SDS), 140 mM NaCl, 1 mM PMSF (all from Sigma-Aldrich)], then incubated for 30 minutes on ice. The lysate was clarified by centrifugation at 13400 ×*g* for 20 minutes at 4°C and the supernatant containing proteins was collected and the concentration determined using the Bicinchoninic Acid (BCA) protein assay (Sigma-Aldrich).

### HISTONE PROTEIN EXTRACTION

Histone proteins were extracted from BV2 microglia cultured in 6-well plates. Cells were washed with 1 mL PBS then scraped into 1 mL ice-cold PBS and pelleted by centrifugation at 400 ×*g* for 5 minutes at room temperature. The cell pellet was resuspended in 400 μL of Triton Lysis Buffer per 1×10^7^ cells [0.5% v/v Triton X-100, 2 mM PMSF, 0.02% w/v NaN_3_ (all from Sigma-Aldrich) and PBS] and incubated on ice for 10 minutes. Lysed cells were centrifuged at 6600 ×*g* for 10 minutes at 4°C. The pellet was resuspended in half the volume of Triton Lysis Buffer used earlier then centrifuged at 6600 ×*g* for 10 minutes at 4°C. The nuclei pellet was resuspended in 50 μL of 200 mM HCl (Acros Organics) and histone proteins were extracted overnight at 4°C. The samples were centrifuged at 6600 ×*g* for 10 minutes at 4°C and the supernatant containing histone proteins was collected and the protein concentration determined using the Bradford protein assay (Sigma-Aldrich).

### WESTERN BLOTTING

Protein samples (10 μg) were separated by sodium dodecyl sulfate polyacrylamide gel electrophoresis (SDS-PAGE) and wet-transferred onto PVDF membrane at 30V for 30 minutes. The membranes were blocked overnight at 4°C with blocking solution [5% w/v nonfat dried milk powder, 0.1% v/v Tween® 20 (both from Sigma-Aldrich) and PBS] and then incubated with either; Anti-HDAC1 [1:2000, rabbit polyclonal, Abcam], Anti-HDAC2 [1:2000, rabbit polyclonal, Abcam], Anti-HDAC3 [1:2000, rabbit polyclonal, Abcam], Anti-β-actin [1:10,000, mouse monoclonal, Sigma], Anti-Histone H3 [1:1000, mouse monoclonal, Cell Signaling], Anti-acetyl Histone H3 Lysine 9 [1:1000, rabbit polyclonal, Millipore] or Anti-acetyl Histone H4 Pan-lysine [1:10,000, rabbit polyclonal, Millipore] (all dissolved in blocking solution) for 1 hour at room temperature. Membranes were washed with PBS-0.1% v/vTween® 20 and incubated with an appropriate secondary antibody; Goat-Anti-Rabbit IgG-horseradish peroxidase (HRP) linked [1:2000, Cell Signaling] or Goat-Anti-Mouse IgG-HRP linked [1:2000, Cell Signaling] for 1 hour at room temperature followed by washing as before. Membranes were incubated with an enhanced chemiluminescence (ECL) substrate(Amersham) and exposed to photographic film, a Fujifilm LAS-3000 imaging system or acDigit® Scanner (LICOR). The intensity of each band was quantified and normalized to the β-actin loading control.

### RNA EXTRACTION, REVERSE TRANSCRIPTION AND QUANTITATIVE RT-PCR

Total RNA was extracted from BV2 microglia cells using TRI Reagent®(Sigma-Aldrich) as per manufacturer’s instructions, resuspended in Tris-EDTA (TE), pH 7.5 and concentrations determined using a NanoDrop 2000c (Thermo Scientific). RNA (2.5 μg) was primed for reverse transcription at 65°C for 5 minutes with 1.25 μL of Oligo(dT)15 primers (0.5 μg/μL, Promega),1.25 μL of Random primers (0.5 μg/μL, Promega) in a final volume of 32.5 μL. cDNA was synthesised at 37°C for 60 min using M-MLV Reverse Transcriptase, RNase H Minus (200 U/μL and RNasin® Plus, Promega) with 2 mM dNTP (Bioline) in a final reaction volume of 50 μL. Quantitative PCR (qPCR) reactions were carried out in duplicate using a Rotor Gene 6000 PCR Analyzer (Corbett) using SensiMix™ SYBR® & Fluorescein (Bioline). Each reaction comprised of 50 or 100 ng of sample cDNA, 300 nM of each gene primer, in a final volume of 20 μL. Primers used were: IL-6 5’-CCCAACTTCCAATGCTCTCC and 5’-ACATGGGATTCCACAAAC, TNF-α’-TGAACTTCGGGGTGATCG and 5’-GGGCTTGTCACTCGAGTTTT, U6 5’- CCGCTTCGGCAGCACA and 5’-AACGCTTCACGAATTTGCGT, HDAC1 5’- GACCGCAAGTGTGTGG and 5’-GAGCAACATTCCGGATGGTG, HDAC2 5’- CAACAGATCGCGTGATGACC and 5’-CCCTTTCCAGCACCAATATCC, HDAC3 5’-GACGTGCATCGTGCTCCAGT and 5’-ACATTCCCCATGTCCTCGAAT. A RNA control (no reverse transcription) and a no template control were also run. PCR conditions were: 95°C for 10 min followed by 45 cycles of 95°C for 10 s, 60°C for 15 s and 72°C for 30 s. Relative quantitation of transcript levels was performed using the 2^-ΔΔCt^ method (Livak & Schmittgen, 2001) and the house-keeping gene U6. Data are presented either as mean percentage expression of the vehicle control ± standard error of the mean (SEM) or mean percentage expression of LPS +vehicle condition ± SEM. Statistical analysis for each experimental condition vs. the vehicle condition 10 was performed using a one-way repeated measures ANOVA followed by the Dunnet *post hoc* test (Lew, 2007) at the 5% significance level.

### ENZYME LINKED IMMUNOSORBANT ASSAYS (ELISAs)

Cell culture supernatants were taken from BV2 microglia cells cultured in 24-well plates. The culture medium was removed and centrifuged for 30 seconds at 16000 ×*g* to pellet any detached cells then 950 μL was removed for analysis. The concentration of mouse IL-6 protein was determined in triplicate by 96-well plate format ELISAs (Invitrogen) following manufacturer’s instructions. Data is presented as mean protein concentration ± standard deviation (SD). Statistical analysis comparing the absolute absorbance values for each experimental condition vs. the vehicle control +LPS condition was performed using a Student’s unpaired t-test assuming equal variances at the 5% significance level.

### INHIBITION OF PROTEIN SYNTHESIS

Cells were treated to 500 ng/mL LPS and either vehicle control, 1 µM SAHA or 500 nM apicidin, in the presence or absence of 1 µg/mL cycloheximide (CHX) for 3 hours. Protein synthesis was assessed in the cells using a Click-iT® Plus O-propargyl-puromycin (OPP) Protein Synthesis Assay Kit (Molecular Probes, Life Technologies). Briefly, after 2.5 hours of drug treatments, Click-iT® OPP was added directly to each well to give a final concentration of 20 µM. The cells were incubated for a further 30 minutes at 37°C. The culture medium was removed, the cells washed with PBS, followed by fixation with 100 µL of 4% w/v paraformaldehyde (Sigma-Aldrich) for 15 minutes at room temperature. The cells were then permeabilised with 100 µL of 0.5% v/v Triton X-100 (Sigma-Aldrich) for 15 minutes at room temperature, washed twice with PBS and processed for imaging following the manufacturer’s instructions. Imaging of labelled cells was carried out using the IncuCyte™ FLR with a 10× objective lens. Nine non-overlapping images were taken in phase-contrast and green-fluorescence (excitation wavelength of 450-490 nm) per well. RNA was extracted from parallel treated cultures for qRT-PCR analysis as described above.

## RESULTS

### HDAC INHIBITION SUPPRESSES CYTOKINE EXPRESSION IN BV2 MICROGLIA

Previous studies using isolated murine microglia report that non-selective HDAC inhibitors (SAHA, TSA and VPA) suppress LPS induced inflammation as measured by a reduction in LPS induced cytokine expression (tumour necrosis factor-alpha, TNF-α and interleukin-6, IL-6) (Suh et al., 2010; Kannan et al., 2013). The specific HDAC(s) important in microglia and neuroinflammation are still largely unknown though identification of the specific HDAC is a requisite for the development of any targeted therapy. We first tested the response of LPS activated BV2 microglia cells to the classical HDAC inhibitors SAHA and VPA (for a discussion of this model system see; (Bocchini et al., 1992; Horvath et al., 2008; Henn et al., 2009; Gresa-Arribas et al., 2012; Stansley et al., 2012)). Six hours after LPS stimulation, IL-6 and TNF-α mRNA expression was increased by 3384 ± 271% and 50990 ± 5190% respectively (Fig 1A), and after 24 hours IL-6 protein secretion was increased by 5406 ± 439% (Fig 1F). BV2 cells express the class I HDACs, 1, 2 and 3 with highest levels of HDAC1 and lowest levels of HDAC3 (Fig 1B, C). Treatment with the HDAC inhibitors, SAHA and VPA produced an increase in the level of Histone H4 acetylation levels within 1 hr which was stable over a period of 24 hr (Fig 1D) suggesting these inhibitors provide rapid and stable HDAC inhibition. Activation by LPS in the presence of either 1 μM SAHA or 10 mM of VPA, produced a significantly reduced response in IL-6 mRNA expression by 84.1 ± 2.8% (P =0.004) and 89.7 ± 1.6%respectively and TNF-α mRNA expression by 59.7 ± 3.2% and 77.9 ± 2.5% respectively (Fig 1E). Furthermore, SAHA significantly suppressed the LPS induced increase in IL-6 protein secretion by 85.6 ± 2.5% (Fig 1F).

In order to understand the role of specific HDACs in microglia and neuroinflammation, we tested HDAC inhibitors that show some selectivity towards specific HDAC isoforms in vitro. We treated LPS induced cells with 5 μM MS-275 (which has reported selectivity for HDAC 1(Bradner et al., 2010)), 500 nM apicidin (which has reported selectivity for HDAC1, 2 and 3(Bradner et al., 2010)), or 1µM MI-192, (which has reported selectivity for HDAC2 and 3(Boissinot et al., 2012)). Treatment of BV2 cells with these inhibitors showed a rapid and stable increase in acetylated Histone H4 for apicidin but a gradual increase for MS-275 and MI-192 over a 24 hour period (Fig 2A). Quantification of histone acetylation levels showed that apicidin produced a similar rate of increase to SAHA while MS-275 and MI-192 required longer incubation periods to induce high levels of histone acetylation (Fig 2B). MS-275 and MI-192 are members of the benzamide class of HDAC inhibitors which have previously been shown to bind HDACs with a slower association rate compared to hydroxamic acid inhibitors such as SAHA (Lauffer et al., 2013). Consistent with these data, co-treatment of LPS with apicidin was sufficient to reduce the induction of IL-6 and TNF-α expression by 82.7 ± 2.3% and 50.3 ± 4.5% but co-treatment of MS-275 or MI-192 was not, (not shown). Pre-treatment of BV2 cells, to mitigate the slow kinetics of inhibition, with either apicidin, MI-912 or MS-275 for 24 hr prior to LPS stimulation significantly reduced the LPS stimulated expression of IL-6 and TNF-α (Fig 2C). Together these data suggest that HDAC1 and HDAC2/3 contribute to the inflammatory response in microglia.

### KNOCKDOWN OF HDAC1 OR HDAC2 SUPRRESSES CYTOKINE EXPRESSION IN BV2 MICROGLIA

Although the HDAC inhibitors show selectivity, they are not isoform specific, therefore we used an siRNA approach to specifically knockdown HDAC1 and HDAC2 to determine their involvement in the inflammatory response of microglia. We were able to significantly knock-down HDAC1 protein expression by 62.6 ± 4.5% (Fig 3A) and HDAC2 protein expression by 68.8 ± 7.7% (Fig 3A). Knockdown of HDAC1 resulted in an increased expression of HDAC2 with no change in HDAC3 (Fig 3A), while knockdown of HDAC2 did not result in any change of expression of HDACs 1 or 3 (Fig 3A). Following knockdown, we treated cells to 500 ng/mL LPS for 6 hours and assessed the expression of IL-6 and TNF-α mRNA. We found, that cells in which HDAC1 was knocked down, there was no change in the response to LPS compared to control cells (not shown) but cells in which HDAC2 was knocked down showed a reduced induction of IL-6 (by 48.2 ± 13%) and TNF-α (by 22.0 ± 3.6%) expression in response to LPS (Fig 3B). To determine if the increase in HDAC2 expression, as a result of HDAC1 knockdown, was acting as a compensatory mechanism we used HDAC1 siRNA to knockdown HDAC1 in combination with a titrated amount of HDAC2 siRNA to reduce HDAC2 to levels seen in control cells (Fig. 3, HDAC1 + 2). Using this approach we were able to significantly reduce HDAC1 levels by 63.5 ± 2.4%, while maintaining the level of HDAC2 to that seen in control cells (89.7 ± 6.2%, Fig 3A HDAC1 + 2). The expression of HDAC3 was not significantly altered (116 ± 5% of Scr siRNA). Cells in which HDAC1 levels are reduced but HDAC2 levels are unchanged showed a reduced response to LPS with with IL-6 and TNF-α mRNA levels of 34.8 ± 3.0% and 35.7 ± 4.8% respectively compared with control cells (Fig 3B). In summary our data identify HDAC1 and 2 activities as important contributors to the neuroinflammatory response of microglia. Furthermore, they show redundancy in this function with increased HDAC2 levels being compensatory for reduced HDAC1.

### HDAC INHIBITION IS EFFECTIVE IN REDUCING THE INFLAMMATORY RESPONSE IN THE ABSENCE OF NEW PROTEIN SYNTHESIS

The mechanism by which HDAC inhibitors exert their effects is often assumed to involve increase in gene expression – indeed HDAC inhibitors do result in increased histone acetylation (e.g. Fig 1D) and the association of increased acetylation with increased gene expression was first identified nearly 30 years ago (Hebbes et al., 1988). However recent proteomic data has identified in excess of 4000 proteins that are modified by acetylation (Choudhary et al., 2009; Lundby et al., 2012; Liu et al., 2014), a number comparable to targets of phosphorylation, suggesting that acetylation is involved in many more processes than gene regulation alone. To determine if the anti-inflammatory action of HDAC inhibition results from changes in gene expression we blocked new protein synthesis using cycloheximide and tested the effectiveness of HDAC inhibitors to block IL-6 and TNF-α stimulation by LPS. Incubation of BV2 cells with cycloheximde for 1 or 3 hours completely blocked new protein synthesis as measured by O-propargyl-puromycin incorporation and protein synthesis was blocked under all conditions used to quantify gene expression levels (Fig 4A). Continued exposure to cycloheximide for 6 hr led to cell death (not shown) though cells were still healthy after 3 hour exposure. The presence of cycloheximde did not affect the induction of IL-6 mRNA expression by LPS (Fig 4B, compare left two bars) and did not prevent either SAHA or apicidin inhibiting this response (Fig 4B, right two bars). Thus these data indicate that the mechanism by which HDAC inhibition reduces the inflammatory response in microglia is manifest within 3 hours and does not require new protein synthesis. Together this suggests that increased gene expression resulting from enhanced histone acetylation is not important for ability of HDAC inhibitors to reduce microglia activation and future work should am to identify the important molecular targets.

## DISCUSSION

It has been previously shown that inhibitors of HDACs can reduce the inflammatory response in activated microglia (Chen and Greene, 2004; Faraco et al., 2009; Suh et al., 2010; Kannan et al., 2013) and by supressing microglia activation show neuroprotective effects following transient ischaemia *in vivo* (Kim et al., 2007; Sinn et al., 2007; Xuan et al., 2012; Kim and Chuang, 2014). However, the identity of the important HDACs involved has not been uncovered and the mechanism by which HDAC inhibition is beneficial is yet to be elucidated. Here we have shown that the function of both HDAC1 and HDAC2 contribute to the inflammatory response in microglia and that in the absence of HDAC1, increased HDAC2 levels compensate suggesting that these two HDACs show redundancy in this function. Furthermore the effectiveness of HDAC inhibition in the absence of new protein synthesis suggests that the HDACs are promoting the inflammatory response by regulating the acetylation levels of a non-histone protein rather than increasing levels of gene expression as a result of increased histone acetylation.

Microglia are often referred to as the immune cells of the brain and recently, selective inhibition and genetic knockdown of class I HDACs, was shown to reduce the production of cytokines in the inflammatory response of macrophages (Jeong et al., 2014). In macrophages, knockdown of either HDAC1 or 2 resulted in increased expression of the other and only a combined knockdown of HDACs 1, 2 and 3 resulted in reduced inflammatory response to LPS (Jeong et al., 2014). In T-lymphocytes deletion of HDAC1 resulted in an increase in HDAC2 protein levels but deletion of HDAC2 had no effect on HDAC1 (Dovey et al., 2013). Here we show that in microglial cells, HDAC1 is the most highly expressed class I HDAC and knockdown of HDAC1 resulted in a compensatory increase in the levels of HDAC2 (Fig 3A). Likewise, we did not observe any compensatory increase in the levels of HDAC1 protein upon knockdown of HDAC2 (Fig 3A) however this contrasts to observations made using macrophages (Jeong et al., 2014). The mechanisms resulting in a compensatory increase in one HDAC upon loss of another are not known though HDAC1 does regulate its own promoter (Schuettengruber et al., 2003) and may also repress expression of other HDACs. One prediction of such a model would be that chemical inhibition of HDAC activity would also result in such compensatory increase. However, we did not observe any compensatory changes in HDAC expression in cells treated with HDAC inhibitors (not shown), suggesting it is not brought about by loss of HDAC enzyme activity but is potentially a mechanism involving the absence of the protein itself. In the absence of HDAC1 in T-lymophocytes the levels of SIN3 and MTA2 are reduced which may indicate that incomplete co-repressor complexes are turned over quickly (Dovey et al., 2013). This structural, rather than enzymatic, requirement for HDAC1 may underlie the reason that knockdown of HDAC1, but not inhibition results in a compensatory increase in HDAC2. Additionally, compensatory changes in HDAC protein levels have been observed in the absence of changes in mRNA levels, suggesting the mechanism involves enhanced translation or protein stability (Jurkin et al., 2011).

HDAC1 and 2 do not exist in the cell as isolated enzymes but are components of three independent co-repressor complexes; Sin3, NuRD and CoREST (for a review see (Kelly and Cowley, 2013)). Each co-repressor complex contains two molecules of HDAC which may consist of two molecules of HDAC1, two molecules of HDAC2 or one of each. Others have observed that upon a loss of HDAC1, HDAC2 can become incorporated into the Sin3, NuRD and CoREST multi-protein complexes in its place (Dovey et al., 2013). The compensatory effect of HDAC2 in the inflammatory response may be explained by such a mechanism. Following a loss of HDAC1, HDAC2 is upregulated and this HDAC is incorporated into a specific complex in place of HDAC1. This complex, specifically targets a protein (which regulates the inflammatory response) for deacetylation. Regardless of the HDAC composition, be it two molecules of HDAC1, HDAC2 or one of each, the specificity for the substrate to be deacetylated comes from the complex itself rather than the HDACs. This hypothesis would suggest it doesn’t matter which of the two HDAC isoforms is inhibited, an anti-inflammatory effect depends on a reduction in the number and activity of this specific functional multi-protein complex. Similarly, the compensatory effect of HDAC3 when HDAC1 and 2 are both lost in macrophages (Jeong et al., 2014) may be explained by HDAC3 being in a specific complex that targets the same substrate as the complex with either HDAC1 or 2. Further research is now needed to investigate these hypotheses and identify the complexes (and composition of them) that when inhibited is responsible for the suppression of pro-inflammatory mediator expression in BV2 microglia.

What is the important target of HDAC1 and 2 that promotes the inflammatory response? Our data identify that new protein synthesis is not required for the HDAC inhibitor response. Formally, we cannot rule out a transcriptional response involving increased miRNA expression and subsequent down regulation of a protein targeted by the miRNA(s), however the ability of the inhibitors to show effectiveness within 3 hours makes such a mechanism unlikely. HDAC enzymes were originally characterised by their ability to deacetylate histone proteins, however these are not their only target and the acetylome may contain on the order of 4000 proteins (Choudhary et al., 2009; Liu et al., 2014). Additionally, the original idea, that HDAC inhibition leads to increased histone acetylation and increased gene expression is likely too simplistic because as many genes are repressed as are activated upon HDAC inhibition by SAHA (Peart et al., 2005). The specific HDAC target(s) important for the microglial response has not been unequivocally identified though a number of potential target proteins can be implicated based on a correlation of their acetylation with microglial activation. Perhaps the most studied non-histone protein involved in the inflammatory response and regulated by acetylation is the transcription factor NF-κB (Greene and Chen, 2004). Quiescent NF-κB is restricted to the cytoplasm via its inhibitory binding partner IκB but upon cell stimulation becomes dissociated and moves into the nucleus where it activates target gene expression. Initially, it was proposed that deacetylation of NF-κB enhanced its interaction with IκB and removal from the nucleus however NF-κB can be acetylated at multiple sites and more recent data suggests that deacetylation at specific residues can result in activation of a subset of NF-κB targets (Rothgiesser et al., 2010), thus inhibition of HDACs may enhance the level of acetylated NF-κB, reducing its activity. In support of this idea, Furumai et al, 2011 showed that inhibition of HDACs in HeLa cells, with TSA, caused a reduction in the recruitment of NF-κB, and RNA polymerase II to the promoter of IL-8, which in turn caused a reduction in IL-8 expression (Furumai et al., 2011). Another candidate protein is MKP-1, a member of the MAPK inflammatory signalling pathway and negative regulator of the inflammatory response. In macrophages MKP-1 activity is reduced when it is deacetylated by HDAC1, 2 or 3 (Jeong et al., 2014). MKP-1 is not just expressed in macrophages but also in microglia (Eljaschewitsch et al., 2006) making this another attractive candidate for the functional response observed here.

In summary our new data here highlight a role for HDAC1 and 2 in regulating microglia activation and suggest the mechanism by which they do so involves acetylation of proteins other than histones. Future studies should now be aimed toward identifying which proteins are the important targets. Although HDAC inhibitors have been approved clinically in the treatment of some cancers they are not without side effects and a more complete understanding of their mechanism of action would open doors to more specific therapeutic targets.

---

**Figure 1.**
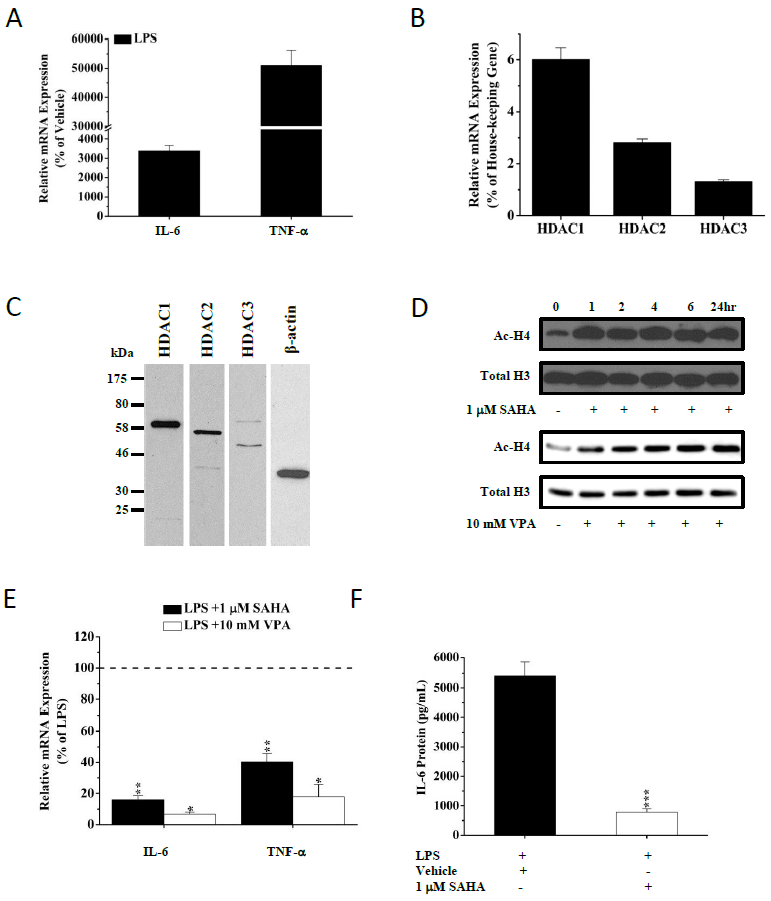
Histone deacetylase (HDAC) inhibition suppress activation of cytokine expression in murine microglia. A) BV2 microglia were stimulated with 500 ng/mL lipopolysacchatide (LPS) and mRNA expression of interleukin-6 (IL-6) and tumor necrosis factor-a (TNF-a) was determined using qPCR. Relative transcript levels were normalized to the U6 gene and expressed as a percentage of the expression in control cells treated with vehicle. Shown are mean ± SEM, *n=3*. B) Quantitative RT-PCR of RNA extracted from BV2 cells. Expression levels expressed as a percentage of the U6 gene. Shown are mean ± SEM, *n=3* C) Western blot analysis of protein extracted from BV2 cells and analysed using anti-HDAC1, HDAC2, HDAC3 and β-actin. Representative blots are shown and positions of molecular weight markers are identified on the left. D) Histone proteins were extracted from control cells and cells treated with 1 μM SAHA or 10 mM VPA for 1, 2, 4, 6, or 24hr. Proteins underwent western blotting for acetylated histone H4 (Ac-H4) and total histone H3. Representative blots are shown. E) BV2 microglia were treated with 500 ng/mL LPS ± 1 mM SAHA or 5 mM valproic acid (VPA) for 6 hours and the mRNA expression of IL-6 and TNF-a was determined using qPCR, normalized to the U6 house-keeping gene and expressed as a percentage of the expression in control cells treated LPS and vehicle shown are mean +/- sem, *n=3*. F) BV2 microglia were treated with 500 ng/mL LPS ± 1 mM SAHA for 24 hours and the changes in IL-6 protein secretion was determined by ELISA. Shown are mean protein concentration ± SEM, *n=3.*). *P < 0.05, **P < 0.01, ***P < 0.001.

**Figure 2.**
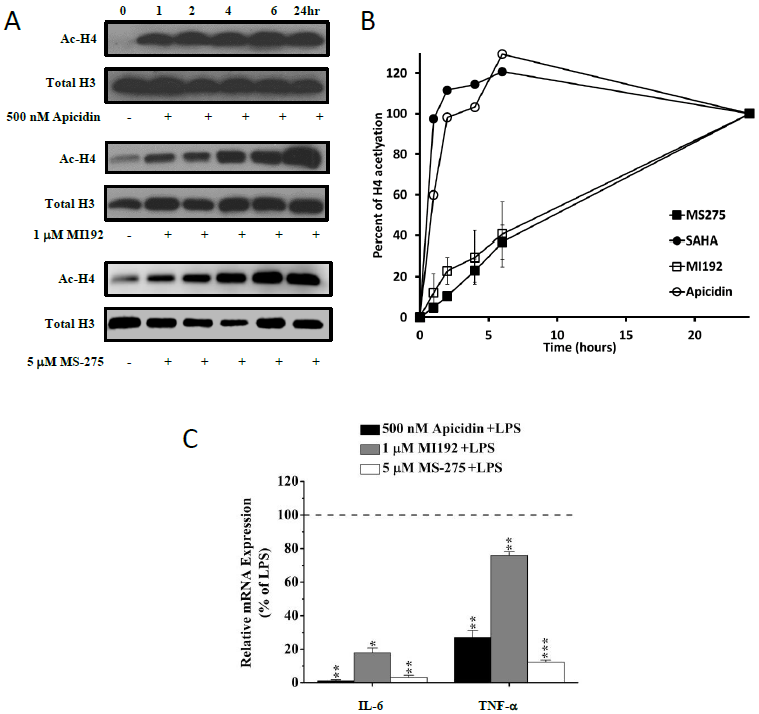
Selective Histone deacetylase (HDAC) inhibitors suppress activation of cytokine expression in murine microglia. A) Histone proteins were extracted from control cells and cells treated with 500nM mM Apicidin, 1μM MI-192 or 5μM MS-275 for 1, 2, 4, 6, or 24hr. Proteins underwent western blotting for acetylated histone H4 (Ac-H4) and total histone H3. Representative blots are shown. B) Quantification of acetylated histone H4 levels, normalised to Histone H3 and expressed relative to acetylation levels at 24hr. C) BV2 microglia were pretreated with HDACi for 24 hours and then stimulated with 500 ng/mL lipopolysacchatide (LPS) for 6 hours. mRNA expression of interleukin-6 (IL-6) and tumor necrosis factor-a (TNF-a) was determined using qPCR. Relative transcript levels were normalized to the U6 house-keeping gene and expressed as a percentage of the expression in control cells treated with vehicle. Shown are mean ± SEM, *3*. *P <0.05,**P < 0.01, ***P < 0.001 compared to LPS stimulated cells.

**Figure 3.**
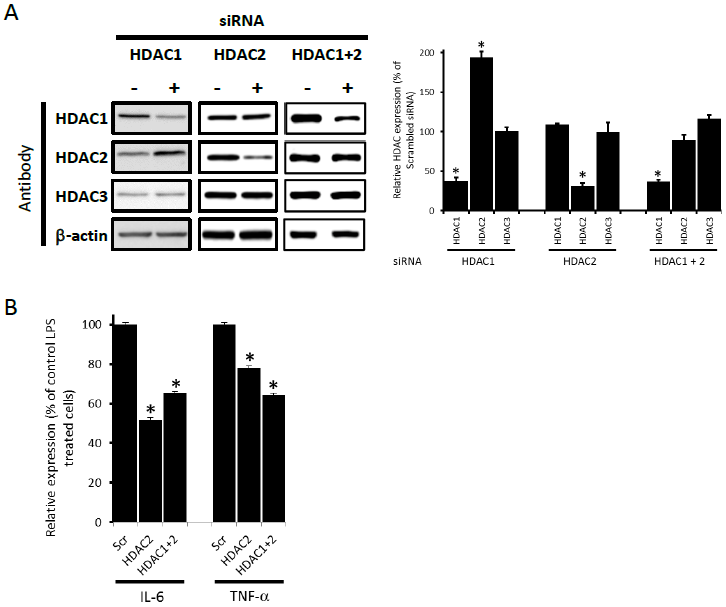
HDAC1 and HDAC2 are involved in the inflammatory response in microglia. A) BV2 microglia were transfected with either scrambled (Scr) siRNA or HDAC1, HDAC2 or HDACs1 and 2 siRNAs. Levels of HDAC1, 2 and 3 and Beta-actin were quantified by immunoblotting. Shown are a representative blots (left) and quantification of protein levels (right) expressed relative to levels in cells treated with Scrambled siRNA. Shown are mean ± SEM, n=3 *p<0.05 compared to scrambled siRNA. B) BV2 microglia transfected with Scr, HDAC1, HDAC2 or HDAC1 and 2 siRNAs were stimulated with LPS and changes in IL-6 and TNF-α mRNA expression was determined using qPCR. Relative transcript levels in each treatment were normalized to the U6 gene and expressed relative to the expression in the Scr siRNA +LPS. Shown are mean ± SEM, n=3. *P < 0.05

**Figure 4.**
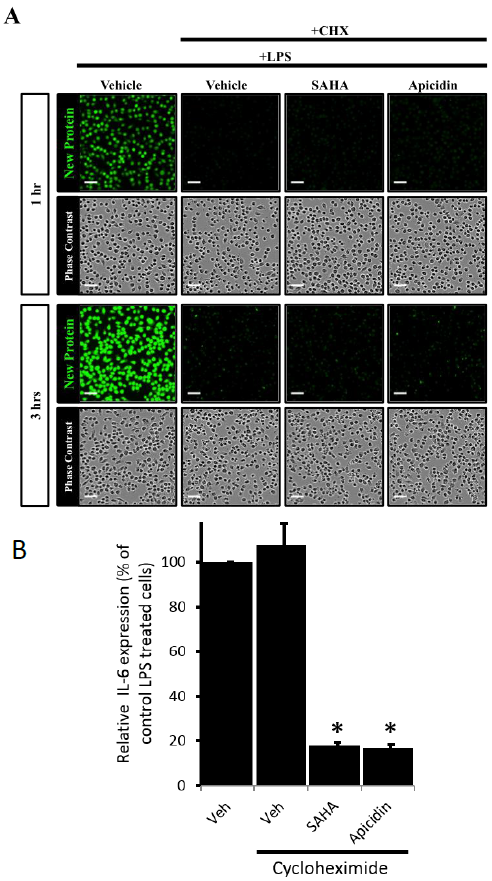
Protein synthesis is not required for HDAC inhibitor efficacy. A) BV2 microgliawere treated for 1 or 3 hours with or without cyclohexamide and incorporation of O-propargyl-puromycin (green, New protein) identified new protein synthesis. Bottom shows phase contrast images, scale bar 50 mm. B) Expression of the IL-6 was measured by quantitative PCR in cells exposed to LPS and treated with vehicle, SAHA or apicidin in the presence of cyclohexamide. Transcript levels for each treatment were normalised to U6 and data shown are mean mRNA expression levels expressed as a percentage of the expression in LPS +vehicle ± SEM, *n*=3,***P<0.001 vs. LPS +vehicle

## References

BeckersT, BurkhardtC, WielandH, GimmnichP, CiossekT, MaierT, SandersK (2007) Distinct pharmacological properties of second generation HDAC inhibitors with the benzamide or hydroxamate head group. International journal of cancer Journal international du cancer 121:1138–1148.

BlockML, ZeccaL, HongJS (2007) Microglia-mediated neurotoxicity: uncovering the molecular mechanisms. Nature reviews Neuroscience 8:57–69.

BocchiniV, MazzollaR, BarluzziR, BlasiE, SickP, KettenmannH (1992) An Immortalized cell line expresses properties of activated microglial cells. Journal of Neuroscience research 31:616–621.

BoyesJ, ByfieldP, NakataniY, OgryzkoV (1998) Regulation of activity of the Transcription factor GATA-1 by acetylation. Nature 396:594–598.

BradnerJE, WestN, GrachanML, GreenbergEF, HaggartySJ, WarnowT, MazitschekR (2010) Chemical phylogenetics of histone deacetylases. Nat Chem Biol 6:238–243.

CameloS, IglesiasAH, HwangD, DueB, RyuH, SmithK, GraySG, ImitolaJ, Duran G, AssafB, LangleyB, KhourySJ, StephanopoulosG, De GirolamiU, RatanRR, FerranteRJ, DangondF (2005) Transcriptional therapy with the histone eacetylase inhibitor trichostatin A ameliorates experimental autoimmune encephalomyelitis. Journal of neuroimmunology 164:10–21.

ChenLF, GreeneWC (2004) Shaping the nuclear action of NF-kappaB. Nat Rev Mol Cell Biol 4525:392–401.

ChenPS, WangCC, BortnerCD, PengGS, WuX, PangH, LuRB, GeanPW, ChuangDM, HongJS (2007) Valproic acid and other histone deacetylase inhibitors induce microglial apoptosis and attenuate lipopolysaccharide-induced dopaminergic neurotoxicity. Neuroscience 149:203–212.

ChoudharyC, KumarC, GnadF, NielsenML, RehmanM, WaltherTC, OlsenJV, MannM (2009) Lysine acetylation targets protein complexes and co-regulates major cellular functions. Science325:834–840.

DoveyOM, FosterCT, ConteN, EdwardsSA, EdwardsJM, SinghR, VassiliouG, BradleyA, CowleySM (2013) Histone deacetylase 1 and 2 are essential for normal T-cell development and genomic stability in mice. Blood 121:1335–1344.

EljaschewitschE, WittingA, MawrinC, LeeT, SchmidtPM, WolfS, HoertnaglH, RaineCS, Schneider-StockR, NitschR, UllrichO (2006) The endocannabinoid anandamide protects neurons during CNS inflammation by induction of MKP-1 in microglial cells. Neuron 49:67–79.

FaracoG, PittelliM, CavoneL, FossatiS, PorcuM, MascagniP, FossatiG, MoroniF, ChiarugiA (2009) Histone deacetylase (HDAC) inhibitors reduce the glial inflammatory response in vitro and in vivo. Neurobiology of disease36:269–279.

FurumaiR, ItoA, OgawaK, MaedaS, SaitoA, NishinoN, HorinouchiS, YoshidaM (2011) Histone deacetylase inhibitors block nuclear factor-kappaB-dependent transcription by interfering with RNA polymerase II recruitment. Cancer Sci 102:1081–1087.

GlassCK, SaijoK, WinnerB, MarchettoMC, GageFH (2010) Mechanisms Underlying inflammation in neurodegeneration. Cell140:918–934.

GlozakMA, SenguptaN, ZhangX, SetoE (2005) Acetylation and deacetylation of non Histone proteins. Gene363:15–23.

GreeneWC, ChenLF (2004) Regulation of NF-kappaB action by reversible acetylation. Novartis Found Symp259:208–217; discussion 218-225.

Gresa-ArribasN, VieitezC, DentesanoG, SerratosaJ, SauraJ, SolaC (2012) Modelling neuroinflammation in vitro: a tool to test the potential neuroprotective effect of anti-inflammatory agents. PloS one7:e45227.

GuW, RoederRG (1997) Activation of p53 Sequence-Specific DNA Binding by Acetylation of the p53 C-Terminal Domain. Cell90:595–606.

HebbesTR, ThorneAW, Crane-RobinsonC (1988) A direct link between core Histone acetylation and transcriptionally active chromatin. EMBO J7:1395–1402.

HennA, LundS, HedtjarnM, SchrattenholzA, PorzgenP, LeistM (2009) The suitability of BV2 cells as alternative model system for primary microglia cultures or for animal experiments examining brain inflammation. Altex26:83–94.

HorvathRJ, Nutile-McMenemyN, AlkaitisMS, DeleoJA (2008) Differential migration, LPS-490 induced cytokine, chemokine, and NO expression in immortalized BV-2 and HAPI cell lines and primary microglial cultures. Journal of neurochemistry107:557–569.

HuE, DulE, SungCM, ChenZ, KirkpatrickR, ZhangGF, JohansonK, LiuR, LagoA, HofmannG, MacarronR, de los FrailesM, PerezP, KrawiecJ, WinklerJ, JayeM (2003) Identification of novel isoform-selective inhibitors within class I Histone deacetylases. J Pharmacol Exp Ther307:720–728.

JeongY, DuR, ZhuX, YinS, WangJ, CuiH, CaoW, LowensteinCJ (2014) Histone deacetylase isoforms regulate innate immune responses by deacetylating mitogen-498activated protein kinase phosphatase-1. Journal of leukocyte biology95:651–659.

JurkinJ, ZupkovitzG, LaggerS, GrausenburgerR, HagelkruysA, KennerL, SeiserC (2011) Distinct and redundant functions of histone deacetylases HDAC1 and HDAC2 in proliferation and tumorigenesis. Cell Cycle10:406–412.

KannanV, BrouwerN, HanischUK, RegenT, EggenBJ, BoddekeHW (2013) Istone deacetylase inhibitors suppress immune activation in primary mouse microglia. Journal of neuroscience research.

KellyRD, CowleySM (2013) The physiological roles of histone deacetylase (HDAC) 1 and 2: complex co-stars with multiple leading parts. Biochem Soc Trans41:741–749.

KhanN, JeffersM, KumarS, HackettC, BoldogF, KhramtsovN, QianX, MillsE, BerghsSC, CareyN, FinnPW, CollinsLS, TumberA, RitchieJW, JensenPB, LichensteinHS, SehestedM (2008) Determination of the class and isoform selectivity of small-molecule histone deacetylase inhibitors. Biochem J409:581

KimHJ, ChuangDM (2014) HDAC inhibitors mitigate ischemia-induced Oligodendrocyte damage: potential roles of oligodendrogenesis, VEGF, and anti-inflammation. American journal of translational research 6:206–223.

KimHJ, RoweM, RenM, HongJS, ChenPS, ChuangDM (2007) Histone Deacetylase inhibitors exhibit anti-inflammatory and neuroprotective effects in a rat Permanent ischemic model of stroke: multiple mechanisms of action. J Pharmacol Exp Ther 517321:892–901.

LaufferBE, MintzerR, FongR, MukundS, TamC, ZilberleybI, FlickeB, RitscherA, FedorowiczG, ValleroR, OrtwineDF, GunznerJ, ModrusanZ, NeumannL, KothCM, LupardusPJ, KaminkerJS, HeiseCE, SteinerP (2013) Histone deacetylase (HDAC) inhibitor kinetic rate constants correlate with cellular histone acetylation but not transcription and cell viability. The Journal of biological chemistry.

LewM (2007) Good statistical practice in pharmacology. Problem 2. British journal of pharmacology 152:299–303.

LiuZ, WangY, GaoT, PanZ, ChengH, YangQ, ChengZ, GuoA, RenJ, XueY, (2014) CPLM: a database of protein lysine modifications. Nucleic Acids Res 42:D531–536.

LundbyA, LageK, WeinertBT, Bekker-JensenDB, SecherA, SkovgaardT, KelstrupCD, DmytriyevA, ChoudharyC, LundbyC, OlsenJV (2012) Proteomic analysis of Lysine acetylation sites in rat tissues reveals organ specificity and subcellular patterns. Cell 530 Rep 2:419–431.

PeartMJ, SmythGKvan LaarRK, BowtellDD, RichonVM, MarksPA, HollowayAJ, JohnstoneRW (2005) Identification and functional significance of genes regulated by structurally different histone deacetylase inhibitors. Proc Natl Acad Sci U S A 102:3697–3702.

PengGS, LiG, TzengNS, ChenPS, ChuangDM, HsuYD, YangS, HongJS (2005) Valproate pretreatment protects dopaminergic neurons from LPS-induced eurotoxicity in rat primary midbrain cultures: role of microglia. Brain research Molecular brain research 134:162–169.

RothgiesserKM, FeyM, HottigerMO (2010) Acetylation of p65 at lysine 314 is important for 540 late NF-kappaB-dependent gene expression. BMC Genomics 11:22.

SchuettengruberB, SimboeckE, KhierH, SeiserC (2003) Autoregulation of mouse histone deacetylase 1 expression. Mol Cell Biol23:6993–7004.

SheinNA, ShohamiE (2011) Histone deacetylase inhibitors as therapeutic agents for acute central nervous system injuries. Molecular medicine17:448–456.

SimoniniMV, CamargoLM, DongE, MalokuE, VeldicM, CostaE, GuidottiA (2006) The benzamide MS-275 is a potent, long-lasting brain region-selective inhibitor of Histone deacetylases. Proceedings of the National Academy of Sciences of the United States of America 103:1587–1592.

SinnDI, KimSJ, ChuK, JungKH, LeeST, SongEC, KimJM, ParkDK, Kun LeeS, KimM, RohJK (2007) Valproic acid-mediated neuroprotection in intracerebral hemorrhage via histone deacetylase inhibition and transcriptional activation. Neurobiology of disease 55226:464–472.

StansleyB, PostJ, HensleyK (2012) A comparative review of cell culture systems for the study of microglial biology in Alzheimer's disease. Journal of neuroinflammation 5559:115.

Strahl BD, Allis CD (2000) The language of covalent histone modifications. Nature 403:41–45.

SuhHS, ChoiS, KhattarP, ChoiN, LeeSC (2010) Histone deacetylase inhibitors suppress the expression of inflammatory and innate immune response genes in human microglia and astrocytes. J Neuroimmune Pharmacol5:521–532.

SuuronenT, HuuskonenJ, PihlajaR, KyrylenkoS, SalminenA (2003) Regulation of microglial inflammatory response by histone deacetylase inhibitors. Journal of neurochemistry 87:407–416.

XuanA, LongD, LiJ, JiW, HongL, ZhangM, ZhangW (2012) Neuroprotective effects of valproic acid following transient global ischemia in rats. Life sciences 90:463–468.

YaoYL, YangWM (2011) Beyond histone and deacetylase: an overview of ytoplasmic histone deacetylases and their nonhistone substrates. J Biomed Biotechnol 2011:146493.

ZhangB, WestEJ, VanKC, GurkoffGG, ZhouJ, ZhangXM, KozikowskiAP, LyethBG (2008) HDAC inhibitor increases histone H3 acetylation and reduces Icroglia inflammatory response following traumatic brain injury in rats. Brain Research 1226:181–191.

ZhangZY, SchluesenerHJ (2013) Oral Administration of Histone Deacetylase Inhibitor MS-Ameliorates Neuroinflammation and Cerebral Amyloidosis and Improves Behavior in a Mouse Model. Journal of neuropathology and experimental Neurology 72:178–185.

ZhangZY, ZhangZ, SchluesenerHJ (2010) MS-275, an histone deacetylase inhibitor, reducesthe inflammatory reaction in rat experimental autoimmune neuritis. Neuroscience 169:370–377.

